# Zinc-binding motif acts as an oxidative stress sensor to regulate TRPM7 channel activity

**DOI:** 10.1101/2020.09.28.316125

**Authors:** Hana Inoue, Takashi Murayama, Takuya Kobayashi, Masato Konishi, Utako Yokoyama

## Abstract

TRPM7 channel activity is negatively regulated by intracellular Mg^2+^. We previously reported that TRPM7 was inhibited by oxidative stress due to an enhancement of the inhibition by intracellular Mg^2+^. In the present study, we aimed to clarify the precise mechanism underlying the TRPM7 inhibition by oxidative stress induced by hydrogen peroxide (H_2_O_2_). Site-directed mutagenesis on full-length TRPM7 revealed that none of the cysteines other than C1809 and C1813 within the zinc-binding motif of the TRPM7 kinase domain were involved in the H_2_O_2_-induced TRPM7 inhibition. When C1809 or C1813 was mutated, full-length TRPM7 was not expressed on the plasma membrane. We, therefore, developed a novel approach in which the full-length TRPM7 is functionally reconstituted by co-expressing the TRPM7 channel domain (M7cd) and the TRPM7 kinase domain (M7kd) as separate individual proteins in HEK293 cells. When M7cd was expressed alone, the current was inhibited by intracellular Mg^2+^ more strongly than in full-length TRPM7. Co-expression of M7cd and M7kd attenuated the current inhibition by intracellular Mg^2+^, and the current was sensitive to oxidative stress, indicating successful reconstitution of a full-length TRPM7-like current. A similar current reconstitution was observed when M7cd was co-expressed with the kinase inactive mutant M7kd-K1645R. Thus, it is suggested that the kinase activity is not essential for the reconstitution. Co-expression of M7cd and M7kd carrying a mutation at C1809 or C1813 failed to restore the full-length TRPM7-like current. No reconstitution was also observed with M7kd carrying a mutation at H1750 and H1807, which are involved in the zinc-binding motif formation together with C1809 and C1813. These data suggest that the zinc-binding motif is essential for the intracellular Mg^2+^-dependent regulation of the TRPM7 channel activity by M7kd, and the cysteines in the zinc-binding motif might play a role in the oxidative stress response of TRPM7.

## Introduction

Magnesium ions are important for numerous cellular functions including cell cycles, channel regulation, enzyme activity, and energy metabolism (reviewed in (Wolf and Trapani, 2008; Romani, 2011; de Baaij et al., 2015). TRPM7 is a Mg^2+^-permeable nonselective cation channel that contains a serine/threonine protein kinase at its carboxyl (C)-terminus (Bates-Withers et al., 2011; Fleig and Chubanov, 2014). Because the TRPM7 channel is activated upon the reduction of intracellular Mg^2+^ ([Mg^*2+*^]_*i*_) and thereby allows a Mg^2+^ influx, TRPM7 is considered to play a role in cellular Mg^2+^ homeostasis (Paravicini et al., 2012; Chubanov et al., 2018). It has been reported that TRPM7-deficient cells exhibit an impairment of proliferation and a decreased survival rate and that these defects are reversed by culturing the cells in a medium containing high Mg^2+^ (Nadler et al., 2001; Schmitz et al., 2003; Schmitz et al., 2005; Chubanov et al., 2016). Global knockout of TRPM7 in mice results in early embryonic death (Jin et al., 2008; Liu et al., 2011; Jin et al., 2012). This lethality might result from the loss of TRPM7 channel activity but not loss of its kinase activity because kinase-inactive TRPM7 knock-in mice (TRPM7^K1646R/K1646R^), in which TRPM7 current is comparable to that in wild-type mice, develop and live normally with no signs of Mg^2+^ deficiency throughout their lifespan (Kaitsuka et al., 2014). Thus, the TRPM7 channel activity, rather than its kinase activity, is pivotal for Mg^2+^ homeostasis under basal conditions.

It is well established that TRPM7 channel activity is regulated by both extracellular and intracellular Mg^2+^. Extracellular Mg^2+^ decreases the current by producing permeation block (Nadler et al., 2001; Monteilh-Zoller et al., 2003), and intracellular Mg^2+^ inhibits the current in a voltage-independent manner (Nadler et al., 2001; Chokshi et al., 2012a, b). Intracellular Mg^2+^ decreases both the open probability (P_o_) and the unitary conductance (^γ^) of TRPM7 (Chokshi et al., 2012b; Mittermeier et al., 2019). It has been proposed that TRPM7 has two intracellular Mg^2+^ sites, with at least one site in the channel domain (Demeuse et al., 2006; Chokshi et al., 2012a), but the precise structural mechanisms underlying the inhibition by intracellular Mg^2+^ remains to be elucidated.

Although the kinase activity of TRPM7 is not essential for channel activity (Schmitz et al., 2003; Matsushita et al., 2005; Demeuse et al., 2006; Inoue et al., 2014; Kaitsuka et al., 2014), the kinase domain might regulate the channel (Schmitz et al., 2003; Desai et al., 2012; Yu et al., 2013). Whole deletion of the kinase domain (TRPM7-Δkinase) increases the [Mg^*2+*^]_*i*_ sensitivity of the TRPM7 channel (Schmitz et al., 2003; Demeuse et al., 2006; Ryazanova et al., 2010), whereas the kinase inactive point mutant (K1645R) does not overtly affect its [Mg^*2+*^]_*i*_ sensitivity (Matsushita et al., 2005). Consistently, a reduction in the TRPM7 current and a defect in Mg^2+^ homeostasis are observed in TRPM7^+/Δkinase^ but not in TRPM7^K1646R/K1646R^ mice (Ryazanova et al., 2010; Kaitsuka et al., 2014; Romagnani et al., 2017). Thus, it is plausible that the structural interactions between the channel domain and the kinase domain are important for the [Mg^*2+*^]_*i*_-mediated regulation of TRPM7 channel activity.

We have previously reported that oxidative stress induced by hydrogen peroxide (H_2_O_2_) inhibited TRPM7 channel activity by enhancing [Mg^2+^]_*i*_-dependent inhibition (Inoue et al., 2014). Our previous study demonstrated that the cysteine modulator N-methyl maleimide (NMM) inhibited the TRPM7 current in a similar Mg^2+^-dependent fashion to H_2_O_2_, suggesting that cysteines act as an oxidative sensor. However, precise molecular mechanisms underlying the oxidative stress-induced TRPM7 inhibition remain to be clarified. In this study, we examined the oxidative stress sensors that are targeted by H_2_O_2_ in TRPM7. Our data suggest that the zinc-binding motif of the TRPM7 kinase domain is responsible for the interaction between the channel domain and the kinase domain to regulate [Mg^*2+*^]_*i*_ sensitivity. The oxidation of cysteines in the zinc-binding motif under oxidative stress might interfere with the interaction to inhibit the TRPM7 current.

## Materials and methods

### Vector constructions

Amino-terminal streptavidin binding peptide (SBP)-tagged full-length wild-type mouse TRPM7 (SBP-mTRPM7-wt) expression vectors were prepared by inserting nucleotides encoding a SBP tag (MDEKTTGWRGGHVVEGLAGELEQLRARLEHHPQGQREP) after the first methionine into an mTRPM7-wt pcDNA5/FRT/TO vector generated previously (Inoue et al., 2014). A full-length cDNA of human TRPM7 (hTRPM7) (pF1KE3491) was purchased from Kazusa Genome Technologies (Chiba, Japan). The hTRPM7 was cloned into a doxycycline-inducible expression vector pcDNA/FRT/TO (Invitrogen, Carlsbad, CA, USA), and the nucleotides encoding the SBP tag were inserted after the first methionine. Mouse TRPM7 channel domain (M7cd, a.a. 1–1509) was prepared by inserting a stop codon and *XhoI* site after 1509 by polymerase chain reaction (PCR) using a *BstXI-SalI* fragment as the PCR template. The TRPM7 kinase domain (M7kd; a.a.1511–1863) was amplified by PCR using an mTRPM7-wt pcDNA5/FRT/TO vector as the template with the primers 5′-CC*ACGCGT*GCCACC<u>ATG</u>TCTAAAGCAGCTTTGTTACC-3′ (MluI site in italics), which includes the ATG start codon (underlined), and 5′-CC*CTCGAG*CTATAACATCAGACGAACAG-3′ (XhoI site in italics). The PCR product was cloned into a pIRES2-EGFP expression vector (BD Biosciences Clontech Franklin Lakes, NJ, USA) in which the MluI site was inserted at the multi cloning site using the MluI and XhoI sites. The full-length coding sequence of mouse TRPM6 (AY333282) was amplified by PCR using cDNA of mouse kidney at the template with the primers 5′-CC*GTCGAC*ATGCAGGTCAAGAAATCCTG-3′ (SalI site in italics) and 5′-CC*GGATCC*TTAAAGGCGTGTGTGATCTT-3′ (M6 reverse primer; BamHI site in italics) and cloned into pBluescript SK(−) (Stratagene, La Jolla, CA, USA) using the SalI and BamHI sites. The TRPM6 kinase domain (M6kd; a.a.1651–2028) was amplified by PCR using the mTRPM6-wt pBluescript SK(−) vector as the template with the forward primer 5′-CC*ACGCGT*GCCACC<u>ATG</u>AAAATGAAGGAAATCAAG-3′ (MluI site in italics), which includes the ATG start codon (underlined), and the M6 reverse primer, and cloned into the pIRES2-EGFP expression vector using the MluI and BamHI sites. Amino acid substitution mutants were generated by site-directed mutagenesis using a QuickChange site-directed mutagenesis kit (Agilent Technologies, La Jolla, CA, USA). The predicted DNA sequences of all constructs were verified by sequencing.

### Baculovirus production

Baculovirus carrying vesicular stomatitis VSVG (virus G protein) on the virus envelope that effectively infects mammalian cells was produced, as described previously (Uehara et al., 2017). cDNAs for wild-type and mutant M7kd were cloned into the modified pFastBac1 vector (Invitrogen) using the MluI and XhoI sites. Baculovirus was produced in Sf9 cells, according to the manufacturer’s instructions. A P2 virus was used for the experiments.

### Cells

Stable and doxycycline-inducible HEK293 cell lines overexpressing the wild-type, mutant, or channel domain of mTRPM7 were generated as previously described (Inoue et al., 2014) using the Flp-In T-REx system (Invitrogen). Cells were maintained in a growth medium, consisting of DMEM supplemented with 10% FBS, 2 mM Glutamax (Invitrogen), 100 μM hygromycin, 15 μM blasticidin, and penicillin-streptomycin (Invitrogen). To induce protein expression, doxycycline (1 μg/mL) was added to the culture medium the day before the experiments. Experiments were performed 16–30 h post-induction.

### Transient expression of M7kd

HEK293 cells that were stably transfected with SBP-tagged or non-tagged mM7cd were plated into a Ф35 mm dish (1 × 10^6^ cell/dish) and cultured in the growth medium for 24 h. M7kd or its mutants encoded in pIRES2-EGFP vectors (2.5 µg/dish) were transfected using lipofectamine 3000 (Invitrogen), according to the manufacturer’s instructions. After an overnight incubation, cells were re-plated on glass cover slips coated with Matrigel (Corning, Corning, NY, USA) and cultured in the growth medium containing doxycycline (1 μg/mL) for 16–30 h.

For baculovirus-mediated expression, HEK293 cells that stably expressed M7cd were plated into a 12-well plate (1.2× 10^5^ cell/well). The cells were cultured for 15–24 h in 900 µL of growth medium containing doxycycline (1 μg/mL) and 100 µL of baculovirus P2 virus solutions.

HEK293 cells expressing M7kd were identified by EGFP fluorescence under an epifluorescence microscope.

### Immunoblotting

Immunoblotting was performed as previously described (Inoue et al., 2014). Proteins in the HEK293 whole-cell lysates were separated by SDS-PAGE using 3–15% linear gradient gels and electrophoretically transferred onto PVDF membranes. To detect full-length TRPM7, its mutants, and M7cd, rabbit anti-TRPM7 antibody (ACC-047; epitope 1146-1165 of human TRPM7, Alomone Labs, Jerusalem, Israel) was used. Rabbit anti-TRPM7+TRPM6 antibody (ab109438; epitope 1800-C-terminus of human TRPM7, Abcam Biochemicals, Bristol, UK) was used to detect M7kd.

### Immunocytochemistry

Stable HEK 293 cells were plated onto glass coverslips (Φ 13 mm) that were coated with Matrigel, and protein expression was induced by doxycycline (1 µg/mL) for 24 h. Cells were fixed in 4% formaldehyde for 10 min at RT, washed three times with PBS, and permeabilized in 0.2% Triton X-100 (Sigma-Aldrich, St. Louis, MO) in PBS for 10 min at RT. Cells were washed three times with cold PBS and incubated with 1% bovine serum albumin (BSA) in PBS for 10 min and then with goat anti-TRPM7 antibody (ab729; Abcam Biochemicals) for 16 h at 4°C. After three washes with 1% BSA-PBS, cells were incubated with a secondary antibody, FITC-conjugated donkey anti goat IgG (ab6881; Abcam). After three washes with 1% BSA-PBS, coverslips were mounted for confocal microscopic imaging (Fluoview ver. 10; Olympus, Tokyo, Japan).

### Patch-clamp experiments

All experiments were conducted at room temperature (23–25°C). The patch electrodes were prepared from borosilicate glass capillaries and had a resistance of 1.5–2.2 MΩ when filled with a pipette solution (see below). Series resistance (<3 MΩ) was compensated (80%) to minimize voltage errors. Currents were recorded using an Axopatch 200B amplifier (Molecular Devices, Union City, CA, USA) that was coupled to a DigiData 1550 A/D and D/A converter (Molecular Devices). Current signals were filtered at 1 kHz using a four-pole Bessel filter and were digitized at 5 kHz. pClamp 10.6 software (Molecular Devices) was used for the command pulse protocol, data acquisition, and analysis. The time courses of the current were monitored by repetitively (every 10 s) applying a ramp pulse from −100 to +100 mV (1-s duration) from a holding potential of 0 mV. The control bath solution consisted of (mM): 135 NaCl, 5 KCl, 1 MgCl_2_, 1 CaCl_2_, 1.2 NaH_2_PO_4_, 10 HEPES, 2 glucose, and 27 mannitol (pH 7.4 adjusted with NaOH, 315 mOsmol/kgH_2_O). The intracellular (pipette) solutions were as follows (mM): 25 CsCl, 110 CsOH, 110 glutamate, 0.2 EGTA, 10 EDTA, and 5 HEPES (pH 7.3 adjusted with CsOH, 290 mOsmol/kgH_2_O). MgSO_4_ was added to vary the free Mg^2+^ concentration ([Mg^2+^]). For the intracellular solution containing 20.9 μM or 0.2 mM [Mg^2+^], EDTA was replaced with HEDTA. For the intracellular solution containing [Mg^2+^] higher than 0.5 mM, EDTA was eliminated. [Mg^2+^] was calculated using MaxChelator software (https://somapp.ucdmc.ucdavis.edu/pharmacology/bers/maxchelator/webmaxc/webmaxcS.htm). H_2_O_2_ or NMM was applied to the extracellular solution for 4 min, and the current amplitudes were analyzed at the end of the application. The current amplitudes just before the application of H_2_O_2_ or NMM were considered to represent the control.

### Statistical analysis

Data are presented as the mean ± standard error of the mean (SEM) of the observations. Comparisons of two experimental groups were made using the Student’s *t-*test. The time-course of the current was compared using a repeated-measures ANOVA. Data were considered to be significant at *p* < 0.05.

To analyze the concentration-dependent inhibition, data were fitted with a monophasic or biphasic concentration–response curve that was provided by the Origin software fitting subroutine (OriginLab, Northampton, MA, USA). The formula for the monophasic concentration–response curves is as follows:

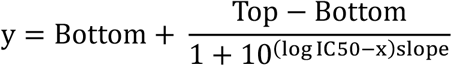

where Bottom and Top are the plateaus at the left and right ends of the curve. IC_50_ is a concentration that gives a half-maximal inhibitory effect and slope is the Hill coefficient.

The formula for the biphasic concentration–response curves with two IC_50_s (IC_50(1)_ and IC_50(2)_, IC_50(1)_ < IC_50(2)_) and two slopes (slope 1 and slope 2) is as follows:

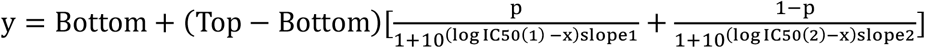

where p is the fraction of high affinity inhibition with IC_50(1)_.

## Results

### Mouse TRPM7 and human TRPM7 were inhibited by H_2_O_2_ in a [Mg^2+^]_i_-dependent manner

It is well established that the TRPM7 current is inhibited by both extracellular and intracellular Mg^2+^. Because the extracellular Mg^2+^ produces voltage-dependent pore block, the inward current is very small in the presence of physiological extracellular divalent cations. Therefore, in our previous studies on overexpressed mouse TRPM7 (mTRPM7) current in HEK293 cells and an endogenous TRPM7 current in mouse white adipocytes, the extracellular divalent cations were eliminated to investigate the effect of oxidative stress that was induced by H_2_O_2_ on both the outward and inward currents (Inoue et al., 2014; Inoue et al., 2019). H_2_O_2_ was shown to inhibit TRPM7 current in a voltage-independent and an intracellular free Mg^2+^ concentration ([Mg^2+^]_*i*_)-dependent manner. To confirm these previous observations, we first recorded a whole-cell current in mTRPM7-wt overexpressing HEK293 cells in the presence of physiological extracellular divalent cations (Figure 1A). The current showed spontaneous activation for approximately 2 min after an establishment of whole-cell configuration (break-in) (Figure 1A). This activation was probably due to a reduction of [Mg^2+^]_*i*_ from approximately 0.9 mM, which was assumed to be in the cytosol, to the [Mg^2+^] in the pipette solutions because the activation rates seemed to be dependent on the [Mg^2+^] in the pipette solutions (Supplemental Figure 1A). The current-voltage (I–V) relationships showed strong outward rectification regardless of the existence of intracellular Mg^2+^ (Figure 1B, C). When 500 μM of H_2_O_2_, which is a concentration that maximally inhibits TRPM7 (Inoue et al., 2014), was applied to the extracellular solution in the presence of 0.2 mM [Mg^2+^]_*i*_, the outward current was markedly inhibited (200.3 ± 28.1 pA/pF and 22.6 ± 3.8 pA/pF at +80 mV, before and 4 min after application of H_2_O_2_, respectively, n = 8) (Figure 1A, B). Although the inward current was very small compared to the outward current due to the existence of extracellular divalent cations, it was also significantly inhibited by H_2_O_2_ (−10.4 ± 1.5 pA/pF and −6.0 ± 1.2 pA/pF at −80 mV, before and 4 min after application of H_2_O_2_, respectively, n = 8). However, the current was completely insensitive to H_2_O_2_ in the absence of intracellular Mg^2+^ (343.0 ± 47.8 pA/pF and 333.4 ± 46.1 pA/pF at +80 mV, −17.9 ± 4.3 pA/pF and −16.1 ± 4.8 pA/pF at −80 mV, before and 4 min after application of H_2_O_2_, respectively, n = 14) (Figure 1A, C). Consistent with previous reports that suggest that TRPM7 has two affinity sites for Mg^2+^ (Chokshi et al., 2012a; Inoue et al., 2014), the TRPM7 current was inhibited by intracellular free Mg^2+^ in a concentration-dependent manner with two IC_50_s (IC_50(1)_ of 5.6 µM and IC_50(2)_ of 558 µM) in the control condition (i.e. before an application of H_2_O_2_). H_2_O_2_ shifted the [Mg^2+^]_*i*_-dependent inhibition curves to lower concentrations with a single IC_50_ of 3.4 µM (Figure 1D). It has been reported that mTRPM7-S1107E mutant is insensitive to [Mg^2+^]_*i*_ (Hofmann et al., 2014; Zhelay et al., 2018). Consistently, the spontaneous current activation after break-in was not observed in mTRPM7-S1107E mutant in whole-cell recordings (Figure 1E). Application of H_2_O_2_ (500 µM) did not affect mTRPM7-S1107E current even in the presence of higher [Mg^2+^]_*i*_ up to 1 mM (Figure 1F). These data were consistent with the concept that a H_2_O_2_-induced decrease in the current is due to an enhancement of [Mg^2+^]_*i*_-dependent inhibition (hereafter referred to as Mg^2+^-inhibition). However, it was also possible that H_2_O_2_ decreased the number of TRPM7 on the plasma membrane by inducing channel internalization. To test this possibility, immunocytochemistry was performed to investigate the membrane expression levels of TRPM7 after the treatment with H_2_O_2_ in mTRPM7-wt overexpressing HEK 293 cells. It was revealed that mTRPM7-wt was clearly detected at the cell boundaries, suggesting the expression on the plasma membrane (Supplemental Figure 1B). A 5-min treatment with H_2_O_2_ (500 μM) did not affect significantly the distribution of TRPM7 on the plasma membrane (Supplemental Figure 1B), suggesting that H_2_O_2_ inhibited the TRPM7 current by reducing channel activity (channel open probability and/or unitary conductance) rather than by channel internalization.

**Figure 1.**
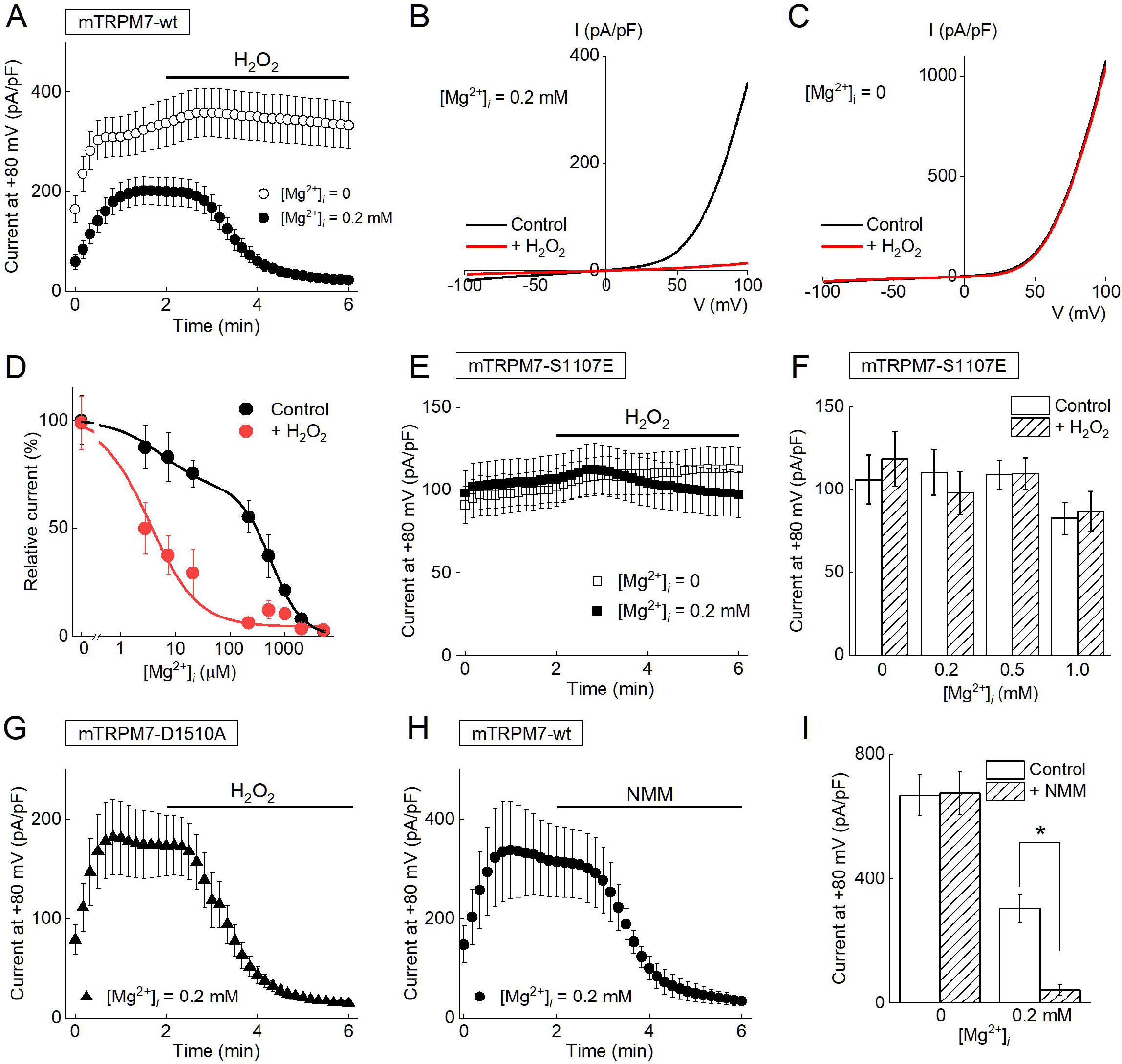
Mouse TRPM7 is inhibited by oxidative stress that is induced by H_2_O_2_ in an [Mg^2+^]_*i*_-dependent manner. (A) Effect of H_2_O_2_ (500 μM) on mTRPM7 current at +80 mV in the absence (open circles) or the presence of 0.2 mM [Mg^2+^]_*i*_ (closed circles). Under the whole-cell clamp mode, ramp command pulses from −100 to +100 V (1-s duration) were applied every 10 s. Each symbol represents the mean ± SEM (vertical bar) of 8–14 recordings. (B) Representative I–V relationship of mTRPM7-wt current recorded in the presence of 0.2 mM [Mg^2+^]_*i*_, before (black line) or 4 min after (red line) the application of H_2_O_2_ (500 μM). (C) Representative I–V relationship of mTRPM7-wt current recorded in the absence of intracellular Mg^2+^, before (black line) or 4 min after (red line) the application of H_2_O_2_ (500 μM). (D) [Mg^2+^]_*i*_-dependent inhibition of mTRPM7 current under control (black circles) and H_2_O_2_-treated (red circles) conditions. The data are presented as a relative current to the mean current that was recorded at 2 min after break-in (i.e. just before application of H_2_O_2_) in the absence of intracellular Mg^2+^. Under control conditions (just before H_2_O_2_ application), the current was inhibited by intracellular free Mg^2+^ with an IC_50(1)_ of 5.6 μM and an IC_50(2)_ of 558 μM (black line). After H_2_O_2_treatment for 4 min, the current was inhibited by intracellular free Mg^2+^ with IC_50_ at 3.4 μM (red line). Each symbol represents the mean ± SEM (vertical bar) of 4–22 observations. (E) A Mg^2+^-insensitive mutant, mTRPM7-S1107E, was not inhibited by H_2_O_2_ (500 μM) in the absence (open squares) or the presence of 0.2 mM [Mg^2+^]_*i*_ (closed squares). Each symbol represents the mean ± SEM (vertical bar) of 13–16 recordings. (F) Mean mTRPM7-S1107E current density did not differ in the absence (open bars) or presence (closed bars) of H_2_O_2_. Each bar represents the mean ± SEM of 8–15 observations. (G) A caspase cleavage-resistant mutant mTRPM7-D1510A was inhibited by H_2_O_2_ (500 μM) in the presence of 0.2 mM [Mg^2+^]_*i*_(closed triangles). Each symbol represents the mean ± SEM (vertical bar) of six recordings. (H) The inhibitory effect of N-methylmaleimide (NMM) (100 μM) on the mTRPM7 current in the presence of 0.2 mM [Mg^2+^]_*i*_ (closed circles). Each symbol represents the mean ± SEM (vertical bar) of five recordings. (I) NMM (100 μM) inhibited mTRPM7 current in the presence of 0.2 mM [Mg^2+^]_*i*_(n = 9), but not in the absence of intracellular Mg^2+^ (n = 4). Each bar represents the mean ± SEM (vertical bar). ^*^*p* < 0.05 by paired *t*-test vs. Control

Furthermore, it was confirmed that human TRPM7 current was inhibited by H_2_O_2_ (500 μM) in a similar [Mg^2+^]_*i*_-dependent fashion (Supplemental Figure 1C, D). These data indicated that mouse and human TRPM7 channel activities are similarly regulated by oxidative stress.

### Cleavage of TRPM7 by caspases was not involved in H_2_O_2_-induced TRPM7 inactivation

It has been reported that TRPM7 is cleaved by caspases at position D1510, and the released TRPM7 kinase domain translocates to the nucleus (Desai et al., 2012; Krapivinsky et al., 2014). The truncated TRPM7 channel domain can be expressed on the plasma membrane (Desai et al., 2012; Duan et al., 2018), although the channel activity is strongly inhibited by intracellular Mg^2+^ (Schmitz et al., 2003). Because H_2_O_2_ activates caspases, it might be possible that caspases mediate the inhibition of TRPM7 by cleaving the kinase domain from TRPM7. To test this possibility, another HEK293 cell line that expressed a caspase cleavage resistant mutant, TRPM7-D1510A, was established (Figure 1G). Similar to TRPM7-wt (Figure 1A), TRPM7-D1510A was markedly inhibited by H_2_O_2_. These data suggest that caspase-mediated cleavage of TRPM7 is not involved in the current inhibition that occurs at least during short (4-min) exposure to H_2_O_2_.

### Screening of cysteines involved in H_2_O_2_-induced TRPM7 inactivation

Since cysteine is one of the most vulnerable amino acid residues to oxidative stress in proteins, it might be possible that H_2_O_2_ oxidizes cysteine(s) in TRPM7 and thereby induces [Mg^2+^]_*i*_-dependent inactivation of the channel activity. Consistent with our previous reports (Inoue et al., 2014), N-methyl maleimide (NMM) (100 µM), a cysteine-modulating reagent, inhibited mTRPM7 and hTRPM7 in a similar manner as that of H_2_O_2_(Figure 1H, I, Supplemental Figure 1E) without affecting localization on the plasma membrane (Supplemental Figure 1B). We then screened for 34 cysteines that are conserved both in mouse and human TRPM7 (Figure 2A) by employing site-directed mutagenesis to identify the oxidation targets during exposure to H_2_O_2_. HEK293 cell lines that stably express each mutant in response to doxycycline treatment were established. The protein expression of each mutant, including a mutant lacking 55 amino (N)-terminus amino acids that contained seven cysteines (ΔNt7C) as well as single- or double-point mutants of cysteine (C) replaced with alanine (A), was confirmed by Western blotting using whole-cell lysate of doxycycline-treated (Dox [+]) HEK 293 cells (Figure 2B). All alanine mutants, except those at the positions C721, C738, C1809, and C1813, expressed robust currents that were significantly larger compered to endogenous currents in doxycycline-untreated (Dox [−]) HEK293 cells in the presence of 0.2 mM [Mg^2+^]_*i*_ (Figure 2C). The current amplitudes in the control conditions were varied among mutants, but H_2_O_2_ consistently decreased the currents to approximately 30% of those before H_2_O_2_ treatment in each mutant, suggesting that these cysteines are not the target for oxidation by H_2_O_2_ to inhibit the current (Figure 2C). Although TRPM7-C721A, -C738A, -C1809A, and -C1813A protein expression was detected in whole-cell lysates by Western blotting (Figure 2B), the current amplitude in these mutants was indistinguishable from that in Dox [−]-cells (Figure 2C).

**Figure 2.**
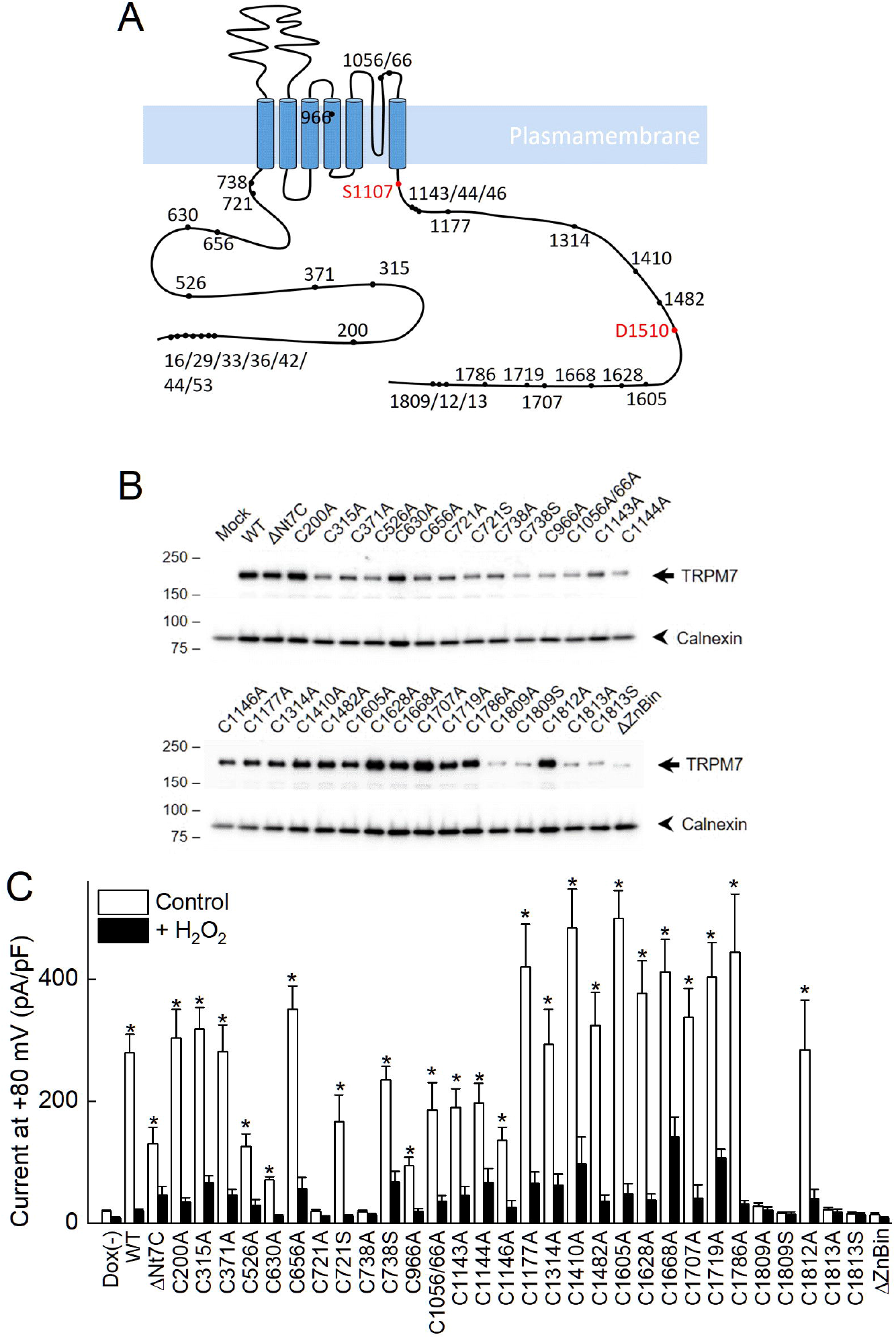
Screening of cysteines in mTRPM7 that were responsible for the oxidative stress-induced inhibition. (A) Schematic diagram of 34 cysteines that are conserved in mouse and human TRPM7. S1107 and D1510 were also depicted (red dots). (B) Expression of TRPM7-wt and its mutants. Immunoblotting of TRPM7 and calnexin (loading control) was performed with HEK293 whole-cell lysates. Numbers on the left represent molecular mass of standards (in kDa). (C) The effect of H_2_O_2_ (500 µM) on mutant TRPM7 current at 0.2 mM [Mg^2+^]_*i*_. Each bar represents the mean ± SEM (vertical bar) of 6–13 observations. ^*^*p* < 0.05 by unpaired *t*-test vs. Dox[−] cells

Consistent with the electrophysiology results, immunocytochemistry revealed that TRPM7-C721A and -C738A seemed to be retained in the intracellular compartments but not expressed on the plasma membrane (Figure 3). Instead of alanine mutants, serine mutants of C721 and C738 could be expressed on the plasma membrane (Figure 3) and exhibit a robust current that was sensitive to H_2_O_2_ (Figure 2C). However, a serine mutant of C1809 and C1813, which are both located in the zinc-binding motif at the C-terminus and are important for structural integrity of TRPM7 kinase domain (Runnels et al., 2001; Yamaguchi et al., 2001), were not expressed on the plasma membrane (Figure 3).

**Figure 3.**
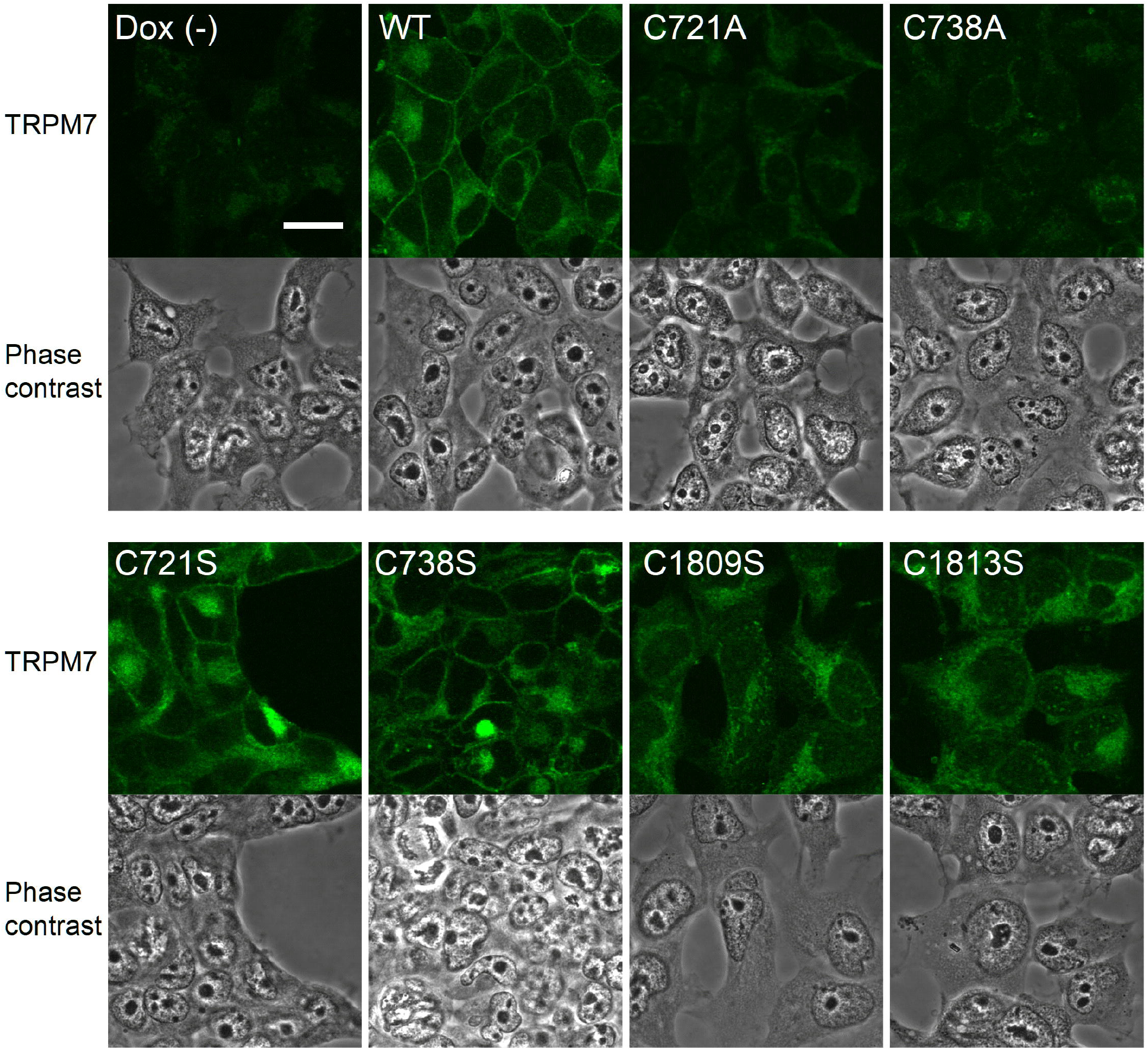
Localization of TRPM7 or its mutants in HEK293 cells. Representative confocal images showing the localization of TRPM7 (green) in doxycycline-untreated (Dox (−)) and mTRPM7-wt (WT), -C721A, -C738A, -C721S, -C738S, -C1809S, and -C1813S-expressing HEK293 cells. Phase contrast images are also shown. Scale bar, 20 μm

### M7cd was functionally expressed in HEK293 cells

Because the cysteine mutations in the zinc-binding motif of TRPM7 interfered with membrane expression, we developed a novel approach in which the full-length TRPM7 was functionally reconstituted by co-expressing M7cd and M7kd as separate proteins in a cell.

We established a HEK293 cell line that stably expresses M7cd (a.a. 1–1510). Whole-cell recordings confirmed the functional expression of M7cd on the plasma membrane in the absence of intracellular Mg^2+^ (Figure 4A, open circles). The current exhibited activation after break-in followed by a rapid rundown in the presence of intracellular Mg^2+^ in M7cd-expressing cells (Figure 4A). Although the rundown of the current continued during 6-min recording, the current amplitude was still dependent on [Mg^2+^]_*i*_ (Figure 4B). Therefore, a concentration–response curve was constructed using the current amplitude at 2 min after break-in (Figure 4C). Consistent with the previous report, which demonstrated that TRPM7-Δkinase (deletion after a.a. 1569) is strongly inhibited by intracellular Mg^*2+*^ (Schmitz et al., 2003), it was revealed that the M7cd current was inhibited by intracellular Mg^2+^ with an IC_50_ of 3.0 µM (Figure 4C). However, M7cd that carries a S1107E mutation (M7cd-S1107E) expressed robust currents constantly in the absence or presence of 0.2 mM [Mg^2+^]_*i*_ (192.5 ± 18.3 pA/pF and 217.9 ± 26.9 pF/pA at +80 mV at 2 min after break-in, in the absence or presence of 0.2 mM [Mg^2+^]_*i*_, n = 10 and 7, respectively) (Figure 4D). These results suggest that the loss of the kinase domain augmented the Mg^2+^-inhibition of TRPM7 channel activity.

**Figure 4.**
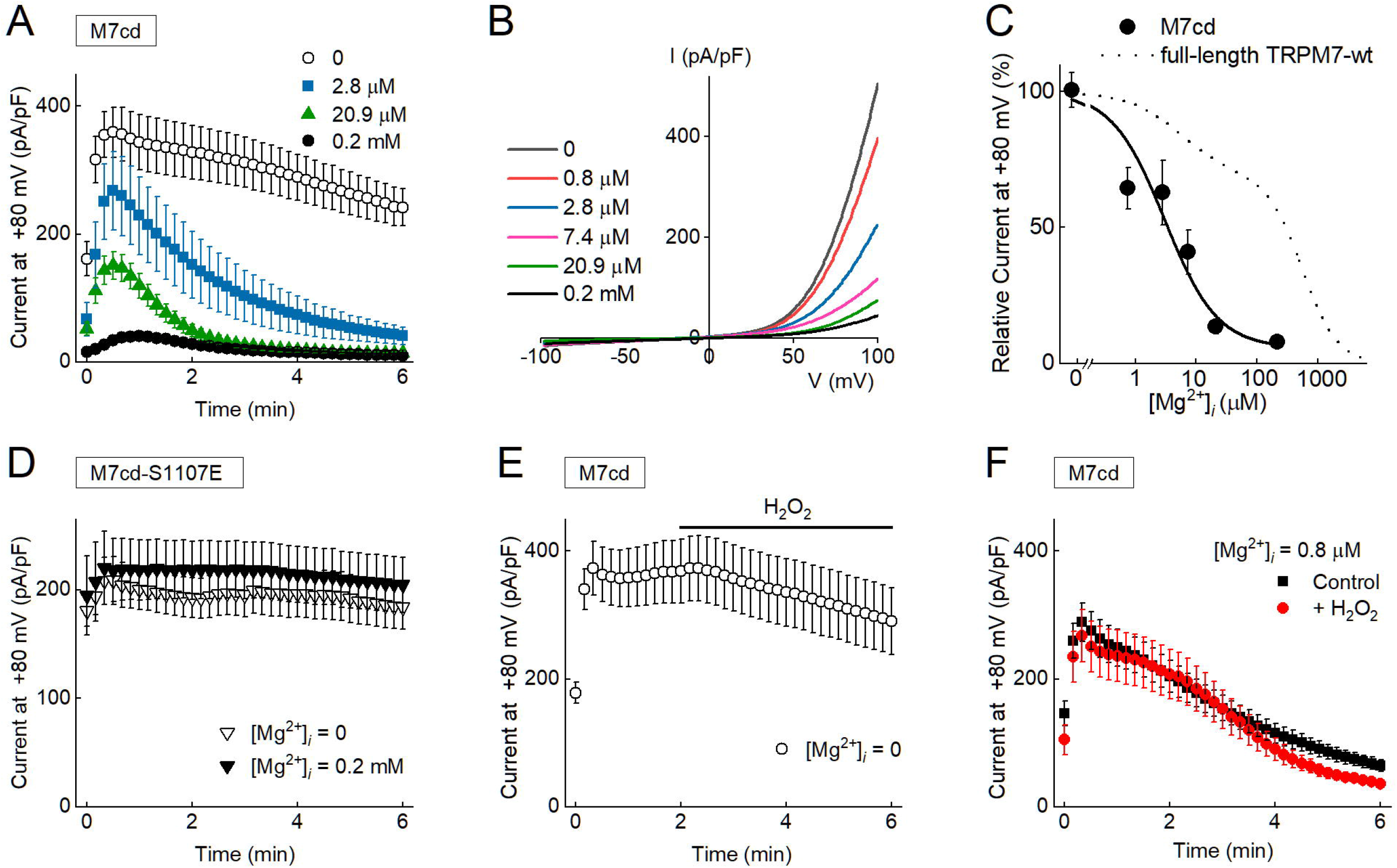
Functional expression of the TRPM7 channel domain (M7cd) in HEK293 cells. (A) Time course of M7cd currents in the absence (open circles) or presence of 2.8 μM (blue squares), 20.9 μM (green triangles), and 0.2 mM [Mg^2+^]_*i*_ (black circles) in HEK293 cells that were treated with doxycycline. Each symbol represents the mean ± SEM (vertical bar) of 9–16 recordings. (B) Representative I–V relationship of M7cd current recorded in the presence of various [Mg^2+^]_*i*_ at 2 min after break-in. (C) [Mg^2+^]_*i*_-dependent inhibition of M7cd current at 2 min. The current was inhibited by intracellular free Mg^2+^ with an IC_50_ of 3.0 μM. Each symbol represents the mean ± SEM (vertical bar) of 14–36 recordings. The dashed line is the [Mg^2+^]_*i*_-dependent curve of the full-length TRPM7-wt current (from Figure 1D). (D) M7cd-S1107E current density was similar in the absence (open inversed triangles) and presence of 0.2 mM [Mg^2+^]_*i*_(closed inversed triangles). Each symbol represents the mean ± SEM (vertical bar) of 7–10 recordings. (E) The effect of H_2_O_2_(500 μM) on M7cd current in the absence of intracellular Mg^2+^ (open circles). Each symbol represents the mean ± SEM (vertical bar) of 19 recordings. (F) The time course of the M7cd current in the presence of 0.8 μM [Mg^2+^]_*i*_ with (red circles) or without (black circles) H_2_O_2_ application at 2 min. Each symbol represents the mean ± SEM (vertical bar) of 17–35 recordings. The current decreased over time even in the control conditions (black circles), and repeated-measures ANOVA revealed that H_2_O_2_ has no significant effect on the current decrease (degrees-of-freedom were corrected using the Greenhouse– Geisser estimate of epsilon, *p* = 0.247).

The effect of H_2_O_2_ on M7cd was tested in the absence or presence of intracellular Mg^2+^ (Figure 4E, F). When we compared with full-length TRPM7-wt (Figure 1A), M7cd current seemed to slightly decrease by an application of H_2_O_2_ (500 μM) in the absence of intracellular Mg^2+^, but there was no statistically significant interaction between the control and H_2_O_2_-treated conditions (Figure 4A, E, repeated-measures ANOVA, *p* = 0.786). Inclusion of Mg^2+^ in the intracellular solution induced rundown even at low concentrations (<1 μM; Figure 4F). H_2_O_2_-treatement did not significantly affect the time-course of current decrease (Figure 4F, *p* = 0.247). Taken together with the results that cysteine mutations in M7cd did not affect the oxidative stress response (Figure 2), these data suggest that M7cd is not the target for H_2_O_2_ to enhance Mg^2+^-inhibition of TRPM7.

### Functional reconstitution of TRPM7 by co-expression of M7cd and M7kd

To test the effect of M7kd co-expression on the M7cd current, M7kd (a.a. 1511–1863) was separately expressed in M7cd-expressing HEK293 cells. In M7cd and M7kd co-expressing cells, the spontaneous current activation after break-in was similar to full-length TRPM7-wt both in the absence and presence of 0.2 mM [Mg^2+^]_*i*_ (Figure 5A). H_2_O_2_ (500 µM) significantly inhibited the reconstituted current in the presence of 0.2 mM [Mg^2+^]_*i*_ (565.7 ± 68.9 pA/pF and 14.0 ± 3.2 pA/pF at +80 mV, before and 4 min after application of H_2_O_2_, n = 8, respectively), but not in the absence of Mg^2+^ (1226.1 ± 240.2 pA/pF and 852.5 ± 184.9 pA/pF at +80 mV, before and 4 min after application of H_2_O_2_, n = 8, respectively) (Figures 5A–C). The concentration–response data of the reconstituted current can be fitted by the formula for a biphasic curve, and this provides an IC_50(1)_of 7.6 μM and IC_50(2)_ of 986 μM (Figure 5D), which were comparable to that of full-length TRPM7-wt (Figure 1D). Similar to full-length TRPM7-wt, a 4-min treatment with H_2_O_2_ (500 µM) shifted the curve leftward, with a single IC_50_of 3.0 µM (Figure 5D). Thus, M7kd attenuates Mg^2+^-inhibition, and reconstitutes a full-length TRPM7-like current that conserves sensitivity to oxidative stress, regardless of whether it is tethered to M7cd.

**Figure 5.**
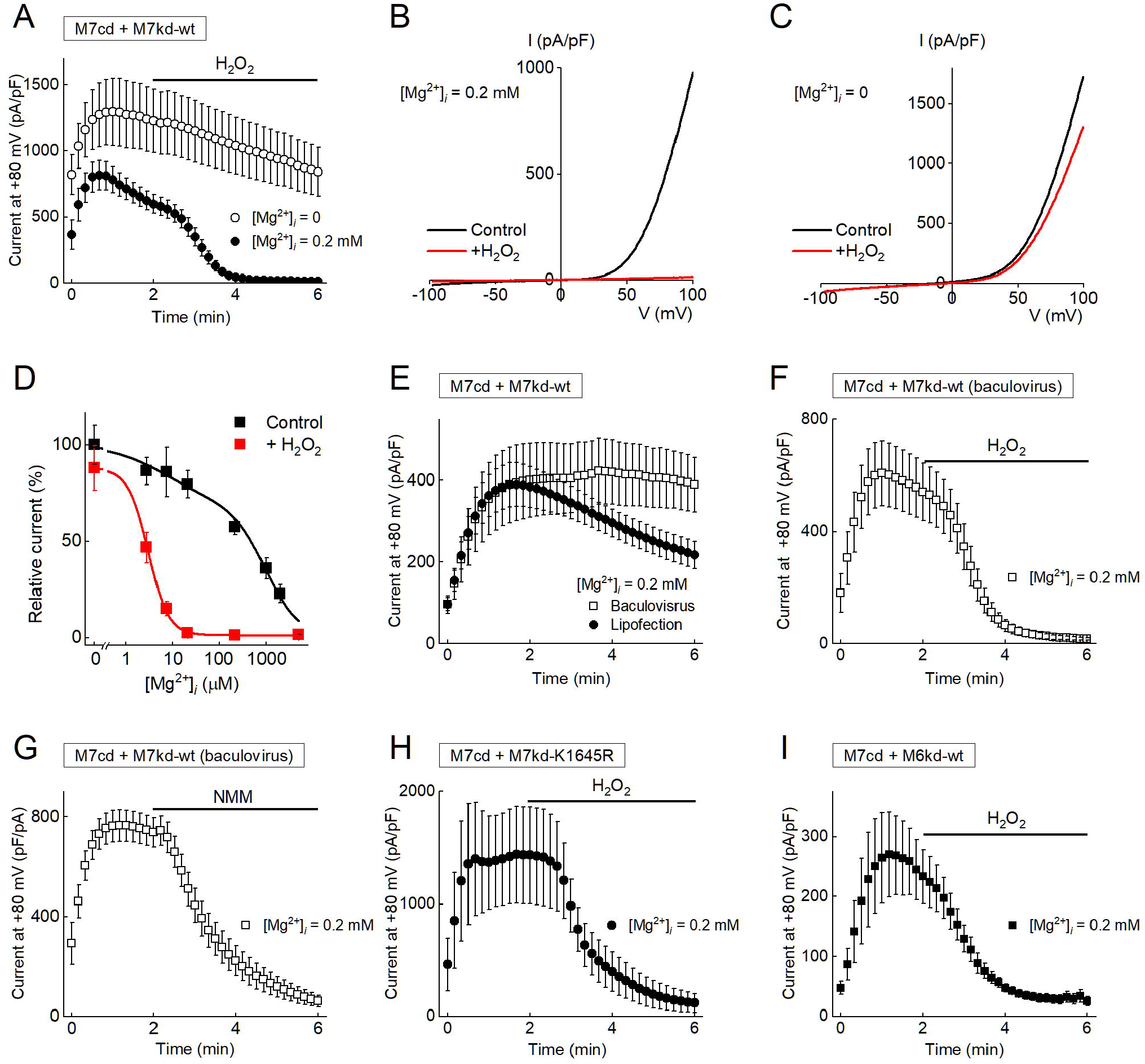
Functional reconstitution of the TRPM7 current by co-expression of M7kd in M7cd-expressing cells. (A) Transient expression of M7kd-wt in M7cd-expressing cells resulted in an increase in the M7cd current density that was inhibited by H_2_O_2_ (500 μM) in the presence of 0.2 mM [Mg^2+^]_*i*_ (closed circles), but not in the absence of intracellular Mg^2+^ (open circles). Each symbol represents the mean ± SEM (vertical bar) of 8–9 recordings. (B, C) Representative I–V relationships of the M7cd current that was recorded in M7cd-expressing cells transfected with M7kd-wt in the presence (B) or absence of 0.2 mM [Mg^2+^]_*i*_ (C), before (black line) or 4 min after (red line) the application of H_2_O_2_ (500 μM). (D) [Mg^2+^]_*i*_-dependent inhibition of M7cd with co-expression of M7kd-wt in the control (black squares) and H_2_O_2_-treated (red squares) conditions. The data fit well with a biphasic concentration–response curve, assuming the fraction of high affinity inhibition of 0.314 estimated for full-length TRPM7-wt (in Figure 1D). Under control conditions, the current was inhibited by intracellular free Mg^2+^ with an IC_50(1)_of 7.6 μM and an IC_50(2)_ of 986 μM. After H_2_O_2_ treatment, the current was inhibited by intracellular free Mg^2+^ with an IC_50_ of 3.0 μM. Each symbol represents the mean ± SEM (vertical bar) of 5–28 observations. (E) The time course of the M7cd current when M7kd-wt was transfected by either lipofection (closed circles) or baculovirus infection (open squares) in the presence of 0.2 mM [Mg^2+^]_*i*_. Each symbol represents the mean ± SEM (vertical bar) of 8–20 observations. (F) The effect of H_2_O_2_ (500 μM) on the M7cd current in the presence of 0.2 mM [Mg^2+^]_*i*_ in cells that were transfected with M7kd-wt by baculovirus infection. Each symbol represents the mean ± SEM (vertical bar) of eight recordings. (G) The effect of NMM (100 μM) on M7cd current in the presence of 0.2 mM [Mg^2+^]_*i*_ in cells that were transfected with M7kd-wt by baculovirus infection. Each symbol represents the mean ± SEM (vertical bar) of six recordings. (H) The effect of co-expression with kinase-inactive mutant M7kd-1645R on M7cd current. Each symbol represents the mean ± SEM (vertical bar) of four recordings. (I) The effect of TRPM6 kinase domain (M6kd) co-expression on M7cd current. Each symbol represents the mean ± SEM (vertical bar) of eight recordings.

During whole-cell recordings in M7cd-expressing HEK293 cells that were transfected with M7kd-expression vector by lipofection, the current sometimes decreased after spontaneous activation before application of H_2_O_2_ (Figures 5A, E, closed circles). Because M7kd was expressed as a cytosolic mobile protein rather than as a plasma membrane-anchored protein, it could be easily diluted with the pipette solutions during whole-cell recordings. Thus, the decrease in the current might result from a decrease in M7kd after break-in. Consistently, when M7kd was expressed with a baculovirus vector that was supposed to enable highly efficient M7kd expression, the current was maintained at least during 6-min recordings (Figure 5E, open squares). It was suggested that the amount of M7kd was sufficient even after dilution with the pipette solution when M7kd was expressed with a baculovirus vector. H_2_O_2_(500 µM) and NMM (100 µM) inhibited the reconstituted current in HEK293 cells that co-express M7cd and M7kd by baculovirus transfection (Figure 5F, G). Therefore, transfection of M7kd with either lipofection or the baculovirus could reconstitute a comparable full-length TRPM7.

### Kinase activity was not involved in the M7kd-mediated regulation of Mg^2+^-inhibition of M7cd

We previously reported that a full-length, kinase-inactive point mutant TRPM7-K1645R is inhibited by H_2_O_2_ in a similar manner as that of TRPM7-wt (Inoue et al., 2014). Co-expression of M7kd-K1645R in M7cd-expressing cells also generated a substantial current that was inhibited by H_2_O_2_ (500 μM) (Figure 5H), indicating reconstitution of the full-length TRPM7-K1645R. It is suggested that the kinase activity is not essential to relieve M7cd from Mg^2+^-inhibition. Consistent results were obtained when M7cd was co-expressed with a kinase domain of TRPM6 (M6kd), which is the closest homologue of TRPM7, but their kinase functions are not redundant (Schmitz et al., 2005) (Figure 5I). M6kd (a.a. 1650–2028) could also attenuate Mg^2+^-inhibition of M7cd, possibly because of their structural similarity such as the presence of a zinc-binding motif at the C-terminus. The reconstituted current was also inhibited by H_2_O_2_ (500 μM) (Figure 5I).

### Mutations of cysteines in the zinc-binding motif interfered with the functional interaction between M7cd and M7kd, which attenuates Mg^2+^-inhibition of TRPM7 current

To examine the effect of mutations of C1809 and C1813, M7kd-C1809S or -C1813S was expressed in M7cd-expressing HEK293 cells. M7cd and M7kd protein expression was confirmed by Western blotting of whole-cell lysates using different antibodies targeting M7cd (ACC-047) and M7kd (ab109438) (Figure 6A). M7kd-wild type (wt) and the mutants M7kd-C1809S and -C1813S were expressed in HEK293 cells that stably expressed M7cd. The functional expression of M7cd on the plasma membrane was also confirmed in cells co-expressing M7cd and mutant M7kd by the whole-cell recordings in the absence of intracellular Mg^2+^ (Figure 6B, C, open circles).

**Figure 6.**
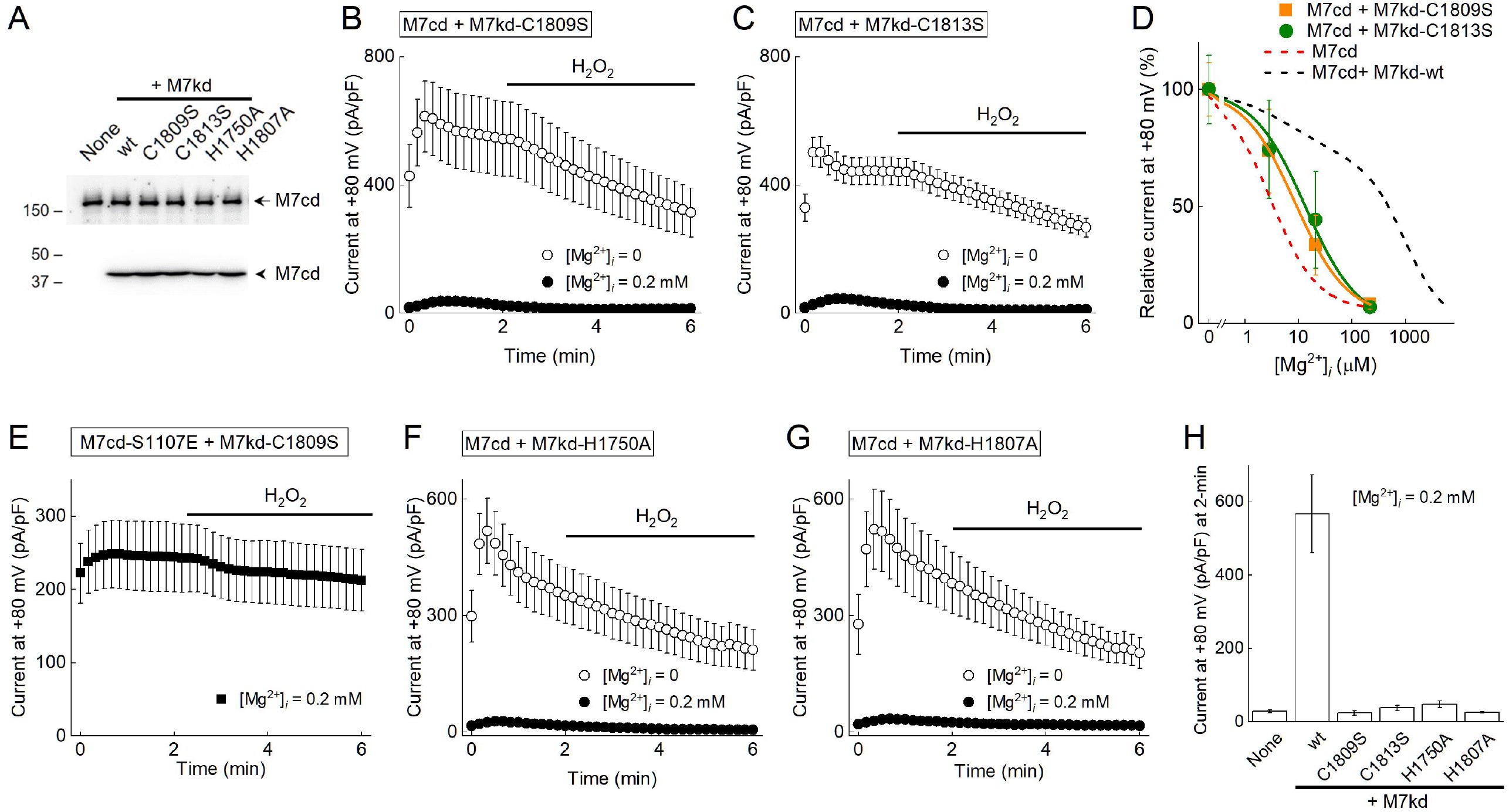
Mutation in the zinc-binding motif in M7kd disrupts the interaction between M7cd and M7kd, which attenuates Mg^2+^-inhibition of the M7cd current. (A) Expression of M7cd and M7kd in M7cd-expressing HEK293 cells transfected with M7kd-wt (lane 2), M7kd-C1809S (lane 3), M7kd-C1813S (lane 4), H1751A (lane 5), and H1808A (lane 6). M7cd expression (172 kDa, arrow) is induced by doxycycline treatment. M7kd expression (40 kDa, arrowhead) is induced by baculovirus infection. (B–C) The effect of M7kd-C1809S (B) or -C1813S (C) expression on M7cd current induced by baculovirus infection in the absence (open circles) or presence of 0.2 mM [Mg^2+^]_*i*_ (closed circles). Each symbol represents the mean ± SEM of 9–17 recordings. (D) [Mg^2+^]_*i*_-dependent curves of M7cd current with co-expression of M7kd-C1809S (orange squares) or -C1813S (green circles) in control conditions. The current was inhibited by intracellular free Mg^2+^ with an IC_50_ of 8.7 μM and 13.4 μM in M7cd-expressing cells with co-expression of M7kd-C1809S and -C1813S, respectively. Each symbol represents the mean ± SEM of 7–17 recordings. The dashed lines are the [Mg^2+^]_*i*_-dependent curve of the M7cd current with (black) or without co-expression of M7kd-wt (red) (from Figures 4C, 5D). (E) M7kd-C1809S expression did not affect the M7cd-S1107E current. Each symbol represents the mean ± SEM of six recordings. (F, G) The effect of M7kd-H1750A (F) or -H1807A (G) expression on M7cd current by baculovirus infection in the absence (open circles) or presence of 0.2 mM [Mg^2+^]_*i*_ (closed circles). Each symbol represents the mean ± SEM of 8–9 recordings. (H) Summary of M7cd current densities at 2 min after break-in in M7kd-wt (wt), -C1809S, - C1813S, -H1750A, or -H1807A-coexpressing cells. Each bar represents the mean ± SEM of 8–9 recordings.

In these settings, co-expression of M7kd-C1809S or -C1813S did not attenuate the Mg^2+^-inhibition of the M7cd current (Figures 6B-D, H). The [Mg^2+^]_*i*_-dependent curves in M7kd-C1809S or -C1813S co-expressing cells were similar to that in cells expressing M7cd alone (IC_50_ of 8.7 for M7kd-C1809S and 13.4 µM for M7kd-C1813S) (Figure 6D). Co-expression of M7kd-C1809S did not affect M7cd-S1107E (Mg^2+^ insensitive mutant) current either before or after an application of H_2_O_2_ (500 µM) (Figure 6E), suggesting that the mutation of C1809 in M7kd does not have a direct inhibitory effect on the current *per se*.

The crystal structure of the TRPM7 kinase domain (a.a. 1521–1863) reveals that a zinc atom is coordinated by H1750, H1807, C1809, and C1813 (Yamaguchi et al., 2001). Therefore, we examined the involvement of H1750 and H1807 in the Mg^2+^-inhibition of the M7cd current (Figures 6F–H). Co-expression of M7kd-H1750A or -H1807A failed to reconstitute the full-length TRPM7, indicating that the structural integrity of M7kd supported by the zinc-binding motif is important for attenuating the Mg^2+^-inhibition of M7cd.

Thus, it is indicated that the zinc-binding motif plays a key role in the functional interaction between M7cd and M7kd in attenuating the Mg^2+^-inhibition of the TRPM7 channel activity. Our data suggest that the oxidation of C1809 or C1813 by H_2_O_2_disrupts the functional interaction between M7cd and M7kd to enhance the Mg^2+^-inhibition of the TRPM7 channel.

## Discussion

In the present study, we found novel mechanisms that support the regulation of TRPM7 channel activity by its kinase domain (Figure 7). It was revealed that intramolecular interactions between the channel domain and the kinase domain increase the TRPM7 current by attenuating TRPM7 inhibition by intracellular Mg^2+^. Mutations of residues in the zinc-binding motif (H1750A, H1807A, C1809, and C1813), which are important for the structural integrity of the TRPM7 kinase domain, diminished the effects of the kinase domain on the TRPM7 current. Our data suggest that oxidative stress inhibits TRPM7 probably via oxidation of C1809 and/or C1813. Oxidation of these cysteines might disrupt the proper structure of the kinase domain and interfere with the interaction between the channel domain and the kinase domain to enhance the inhibition of TRPM7 by intracellular Mg^2+^.

**Figure 7.**
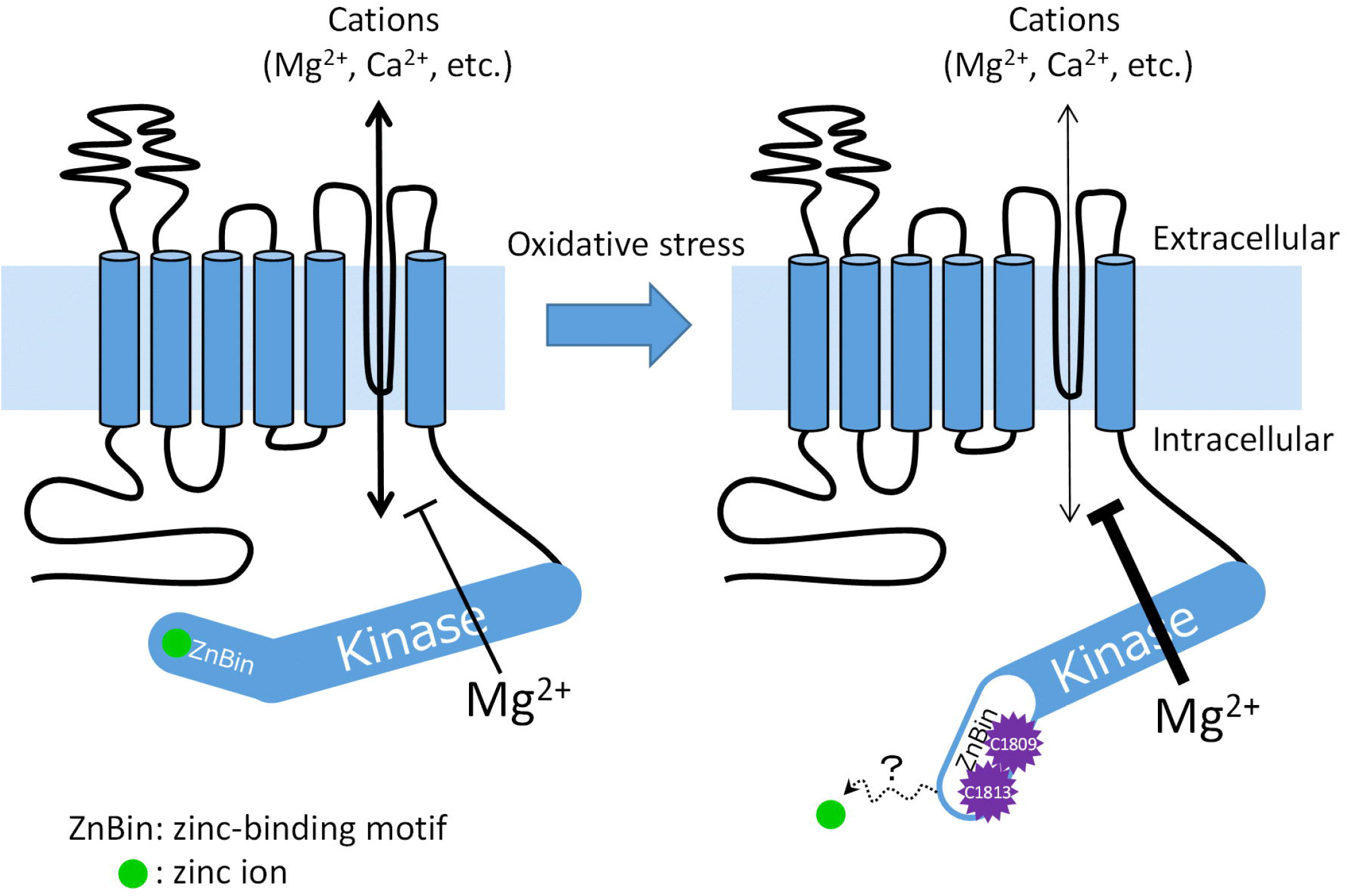
Schematic illustration of TRPM7 regulation by oxidative stress. (Left) Under basal conditions, Mg^2+^-inhibition of TRPM7 current is attenuated by the kinase domain possibly via the interaction with the channel domain. (Right) Oxidative stress causes the oxidation of the cysteines (C1809 and C1813) in the zinc-binding motif, which may disrupt the interaction between the channel domain and the kinase domain and enhance the inhibition of TRPM7 by intracellular Mg^2+^.

Oxidation of cysteines and methionines in proteins regulates numerous redox-sensitive physiological functions (Rhee et al., 2000; Poole and Nelson, 2008; Reddie and Carroll, 2008; Klomsiri et al., 2011). Among TRPM family members, TRPM2 and TRPM6 have been reported to be regulated by oxidative stress via direct oxidation of a certain methionine residue (Cao et al., 2010; Kashio et al., 2012). Our present study suggests that C1809 and C1813 act as an oxidative stress sensor in TRPM7. The crystal structure of the TRPM7 kinase domain has revealed that these cysteines coordinate a zinc ion with H1750 and H1807 (Yamaguchi et al., 2001). This zinc-binding motif is important for the structural integrity of the TRPM7 kinase domain, and thus, mutation of C1809 and C1813 results in a loss of kinase activity (Runnels et al., 2001). Because the kinase-inactive M7kd-K1645R as well as M6kd, which is a functionally non-redundant kinase to M7kd (Schmitz et al., 2005), attenuated the Mg^2+^-inhibition of M7cd current in a manner that was comparable to that of M7kd-wt (Figure 5H, I), the kinase activity is not likely involved in the regulation of [Mg^2+^]_*i*_-sensitivity. However, M7kd-C1809 or -C1813 failed to increase the M7cd current (Figure 6B–D, H). Furthermore, our experiments showed that M7kd-H1750A or -H1807A also failed to attenuate the Mg^2+^-inhibition of the M7cd current (Figure 6F–H). From these results, it is suggested that the structural integrity of M7kd that is guaranteed by the zinc-binding motif is important for the interaction between M7cd and M7kd to regulate the [Mg^2+^]_*i*_-sensitivity of TRPM7. It has been proposed that the zinc-binding motif acts as an oxidative stress sensor in various proteins to regulate cellular functions in response to redox changes (Ilbert et al., 2006; Kroncke and Klotz, 2009). Similarly, the zinc-binding motif of TRPM7 might be an oxidative sensor that regulates the channel activity.

To induce oxidative stress, 500 μM H_2_O_2_ was added to the extracellular solutions in the present study. It might be questionable that such high concentrations can occur *in vivo*. It has been reported that a gradient between extracellular and intracellular H_2_O_2_ concentrations is 390-fold (Lyublinskaya and Antunes, 2019) or 650-fold (Huang and Sikes, 2014), with the lower concentrations in the cytosol. Thus, [H_2_O_2_]_*i*_was estimated to be approximately 0.8–1.3 μM, when 500 μM H_2_O_2_ was applied to the extracellular solution. Such submicromolar intracellular H_2_O_2_ has been shown to mediate the redox signaling under pathological conditions (Sies, 2017). Furthermore, extracellular H_2_O_2_ inhibits TRPM7 with an IC_50_ of 15.9 μM (Inoue et al., 2014), suggesting that H_2_O_2_ might oxidase C1809 and/or C1813 even within the range of physiological concentrations ([H_2_O_2_]_*i*_ = 1–10 up to 100 nM). Thus, regulation of the TRPM7 channel activity by H_2_O_2_ is a mechanism that may occur under both physiological and pathophysiological conditions.

In our electrophysiological experiments, EDTA or HEDTA was included in the pipette solutions to maintain constant [Mg^2+^]_*i*_even under conditions where Mg^2+^ can influx due to channel activity (i.e., in the presence of extracellular Mg^2+^), when the [Mg^2+^]_*i*_was set at <0.5 mM (Materials and Methods). In addition to Mg^2+^, these chelators bind Zn^2+^ with a dissociation constant (*K*_*d*_) of 2.3×10^−14^ M or 6.6×10^−13^ M, respectively (Krezel and Maret, 2016). Therefore, the use of these chelators might unfold the zinc-binding motif of TRPM7 by depleting a zinc ion under the whole-cell configuration. However, we were able to measure a robust TRPM7 current in the presence of the chelators (Figure 1), suggesting that the TRPM7 zinc-binding motif provides a high-affinity binding site for zinc. Similarly, an extracellular application of a membrane-permeable, zinc-specific chelator, TPEN (100 μM, *K*_*d*_ = 6.4×10^−16^ M) did not cause marked TRPM7 inhibition like H_2_O_2_in our preliminary experiments (data not shown). It has been reported using zinc finger peptide models that a well-packed hydrophobic core in the vicinity of a zinc-binding motif increases the binding constant (*K*_*a*_) to 10^13^–10^16^ M and slows down the kinetics of metal exchange (Seneque and Latour, 2010). In the TRPM7 zinc-binding motif, a zinc atom is integrated into the hydrophobic core and is secluded from solvent (Yamaguchi et al., 2001). Thus, the zinc-binding motif of TRPM7 might be folded as intact, even in the presence of the chelators, until H_2_O_2_ is applied.

Our results demonstrated that M7cd and M7kd interact functionally, but the interaction sites of M7cd with M7kd remain to be identified. Although the protein–protein interaction between M7cd and M7kd was tested using an immunoprecipitation technique in our preliminary experiments, M7kd did not co-immunoprecipitate with M7cd probably due to the low affinity between M7cd and M7kd (data not shown). Consistently, electrophysiology data also suggested weak binding between M7cd and M7kd (Figure 5E). When M7kd was overexpressed in M7cd-expressing cells as a separated individual protein, the current was robust for several minutes after break-in, but decreased over time (Figure 5E). This might be due to a decrease in M7kd concentration in the cytosol through intracellular perfusion with the pipette solution. Thus, because of the weak binding of M7cd and M7kd, it is difficult to identify the interaction sites via a general immunoprecipitation technique or a pull-down assay. Fluorescence resonance energy transfer experiments may be an effective approach to confirm the interaction. Alternatively, structural analysis might be more effective for clarifying the detailed mechanism underlying the interaction. To date, the structure of the TRPM7 channel domain (lacking the kinase domain) and that of the TRPM7 kinase domain (lacking the channel domain) have been resolved independently (Yamaguchi et al., 2001; Duan et al., 2018). However, the whole structure of TRPM7 remains to be identified. Thus, further structural study is necessary to identify the interaction sites between M7cd and M7kd that regulate the channel activity.

It has been proposed that TRPM7 has two intracellular Mg^2+^ sites to inhibit its channel activity (Nadler et al., 2001; Monteilh-Zoller et al., 2003; Chokshi et al., 2012a, b). Consistent with this idea, the concentration–response data for the full-length TRPM7-wt in our study were well explained by two binding sites for Mg^2+^, a high affinity site (IC_50(1)_ < 10 μM) and a low affinity site (IC_50(2)_ > 500 μM) (Figure 1D). The present study revealed that M7cd has a single affinity site with an IC_50_ of 3.0 μM that is quite similar to the IC_50(1)_ of full-length TRPM7 (Figure 4C). Thus, it is suggested that the high affinity Mg^2+^ site is located in M7cd, whereas the low affinity site is located in M7kd. From the results that H_2_O_2_ both shifted the [Mg^2+^]_*i*_-dependent inhibition curves to lower concentrations and converted it to a monophasic curve with IC_50_ of 3.4 μM (highly similar to that of M7cd), it seems as if H_2_O_2_ removed M7kd from the full-length TRPM7-wt. Because the cleavage-resistant mutant, TRPM7-D1510A, remained sensitive to oxidative stress (Figure 1G), H_2_O_2_ might completely interfere with the functional interaction between the channel domain and the kinase domain by oxidizing C1809 and/or C1813, rather than physically cleaving it. It can be also speculated that attenuation of Mg^2+^-inhibition by M7kd is not due to reduction of Mg^2+^ binding affinity to the site in M7cd because the full-length TRPM7 retains the high-affinity Mg^2+^ binding. From the fitting, the fraction of Mg^2+^-inhibition that is related to the high-affinity Mg^2+^ binding is estimated to be approximately 30% of maximal Mg^2+^-inhibition in full-length TRPM7 (Figure 1D). Therefore, M7kd might reduce the transduction from the high-affinity Mg^2+^-binding to channel inhibition by approximately 70%. Because M7kd has a low affinity site for Mg^2+^, the remaining 70% of Mg^2+^-inhibition is induced at high concentrations of Mg^2+^. Thus, H_2_O_2_ inhibits TRPM7 via an apparent enhancement of Mg^2+^-inhibition.

Mutation of cysteines in the zinc-binding motif as well as the alanine mutants of C721 and C738 in full-length TRPM7 were not expressed on the plasma membrane, and instead remained at an intracellular compartment (Figure 3). The results suggest that these cysteines might be important for proper folding of TRPM7 that is expressed on the plasma membrane. A heterozygous mutation of C721 to glycine in TRPM7 has been found in a human pedigree with macrothrombocytopenia (Stritt et al., 2016). In contrast to TRPM7-C721A, which was examined in the present study, the TRPM7-C721G mutant is successfully expressed on the plasma membrane though the current is reduced by 85% of that of TRPM7-wt when expressed in HEK293 cells. The serine mutant of C721 (TRPM7-C721S) could be expressed on the plasma membrane (Figure 3), and the current was comparable to that of TRPM7-wt (Figure 2C). Thus, the results vary based on which amino acid is substituted for C721. The structure defined by C721 might be important for expression on the plasma membrane and for TRPM7 channel activity, but it may not be involved in the oxidative stress-induced inhibition of TRPM7.

There has been an increasing amount of interest in investigating the involvement of TRPM7 channel activity in a range of pathologies, such as ischemia in brain (Aarts et al., 2003; Sun et al., 2009; Chen et al., 2015a) and kidney (Meng et al., 2014; Liu and Yang, 2019) as well as in cancer (Guilbert et al., 2009; Rybarczyk et al., 2012; Chen et al., 2015b; Liu et al., 2020). Downregulation of TRPM7 by small interference RNA or TRPM7 inhibitors alleviates ischemic damage in neurons (Chen et al., 2015a; Sun, 2017). Because TRPM7 is permeable to divalent cations including Ca^2+^ and Zn^2+^, it is conceivable that downregulation of TRPM7 protect cells from Ca^2+^- or Zn^2+^-induced toxicity that is associated with ischemia (Inoue et al., 2010; Sun, 2017). In cancer cells, it has been reported that increased activity of TRPM7 might be involved in cancer cell proliferation and migration (Guilbert et al., 2009; Rybarczyk et al., 2012; Chen et al., 2015b). Thus, downregulation of TRPM7 might be a therapeutic target under these pathological conditions. There are several potent inhibitors of TRPM7, such as waixenecin A (Zierler et al., 2011) and NS8593 (Chubanov et al., 2012), but the binding site of these compounds in the TRPM7 molecule remains largely unknown. Both waixenecin A and NS8593 inhibit TRPM7 current in the absence of intracellular Mg^2+^ or its kinase domain, but the inhibitory mechanisms are different from those of oxidative stress. Based on our present findings, compounds that are designed to interrupt the intramolecular interaction between M7cd and M7kd can be specific inhibitors of TRPM7. In conclusion, oxidative stress inhibits TRPM7 channel activity. Cysteine residues of the zinc-binding motif are suggested to act as the oxidative stress sensor, and its oxidation might interfere with the intramolecular interactions between the channel domain and the kinase domain, thereby increasing Mg^2+^-dependent inhibition of the channel activity.

## Acknowledgements

We thank Dr. N. Fukushima for useful discussions. This work was supported in part by JSPS KAKENHI (JP22460302, H.I.; JP17K08549, H.I.; 19H03404, H.I and T.M.), the Platform Project for Supporting Drug Discovery and Life Science Research (Basis for Supporting Innovative Drug Discovery and Life Science Research [BINDS]) from AMED (JP20am0101080, *support number 0743*, T.M.), and the Tokyo Medical University Research Support Program during Life Events (H.I).

## Author contributions

H. Inoue performed the electrophysiological experiments and cell biological experiments. T. Murayama and T. Kobayashi provided experimental tools and performed biochemical experiments. H. Inoue and T. Murayama analyzed the data. H. Inoue, T. Murayama, M. Konishi, and U. Yokoyama wrote the manuscript. All authors discussed the results and approved the final version of the manuscript.

## Figure legends

*Supplemental Figure 1*

(A) Time course of whole-cell currents in the presence of various [Mg^2+^]_*i*_in full-length mTRPM7-overexpressing HEK293 cells. Each symbol represents the mean ± SEM of recordings at [Mg^2+^]_*i*_, which are shown on the right. The current at 2 min (control, the black vertical dashed line) and 6 min (H_2_O_2,_ the red vertical dashed line) relative to the mean current at 2 min in the absence of Mg^2+^ are plotted in Figure 1D. Each symbol represents the mean ± SEM (vertical bar) of 4–22 observations. (B) Localization of TRPM7 in full-length mTRPM7-wt-overexpressing HEK293 cells. Control conditions (left), 5-min treatment with H_2_O_2_ (500 μM, middle) or NMM (100 μM, right). Confocal images showing the localization of TRPM7 (green) and nuclei (blue). Scale bar, 20 μm. (C) The effect of oxidative stress induced by H_2_O_2_ on the human TRPM7 current. Representative traces of whole-cell currents showing the time course of the current inhibition by H_2_O_2_ (500 μM) in the absence (open circles) or presence (closed circles) of 0.2 mM [Mg^2+^]_*i*_ in human TRPM7 (hTRPM7)-expressing HEK293 cells. (D) H_2_O_2_ (500 μM) inhibited the hTRPM7 current in the presence of 0.2 mM [Mg^2+^]_*i*_ (n = 6), but not in the absence of intracellular Mg^2+^ (n = 5). Each bar represents the mean ± SEM (vertical bar). ^*^*p* < 0.05 vs. Control. (E) NMM (100 μM) inhibited the hTRPM7 current in the presence of 0.2 mM [Mg^2+^]_*i*_ (n = 5), but not in the absence of intracellular Mg^2+^ (n = 4). Each bar represents the mean ± SEM (vertical bar). ^*^ *p* < 0.05 vs. Control

