## Supplementary figures and images for "Zinc-binding motif acts as an oxidative stress sensor to regulate TRPM7 channel activity"

### Supplemental Figure 1

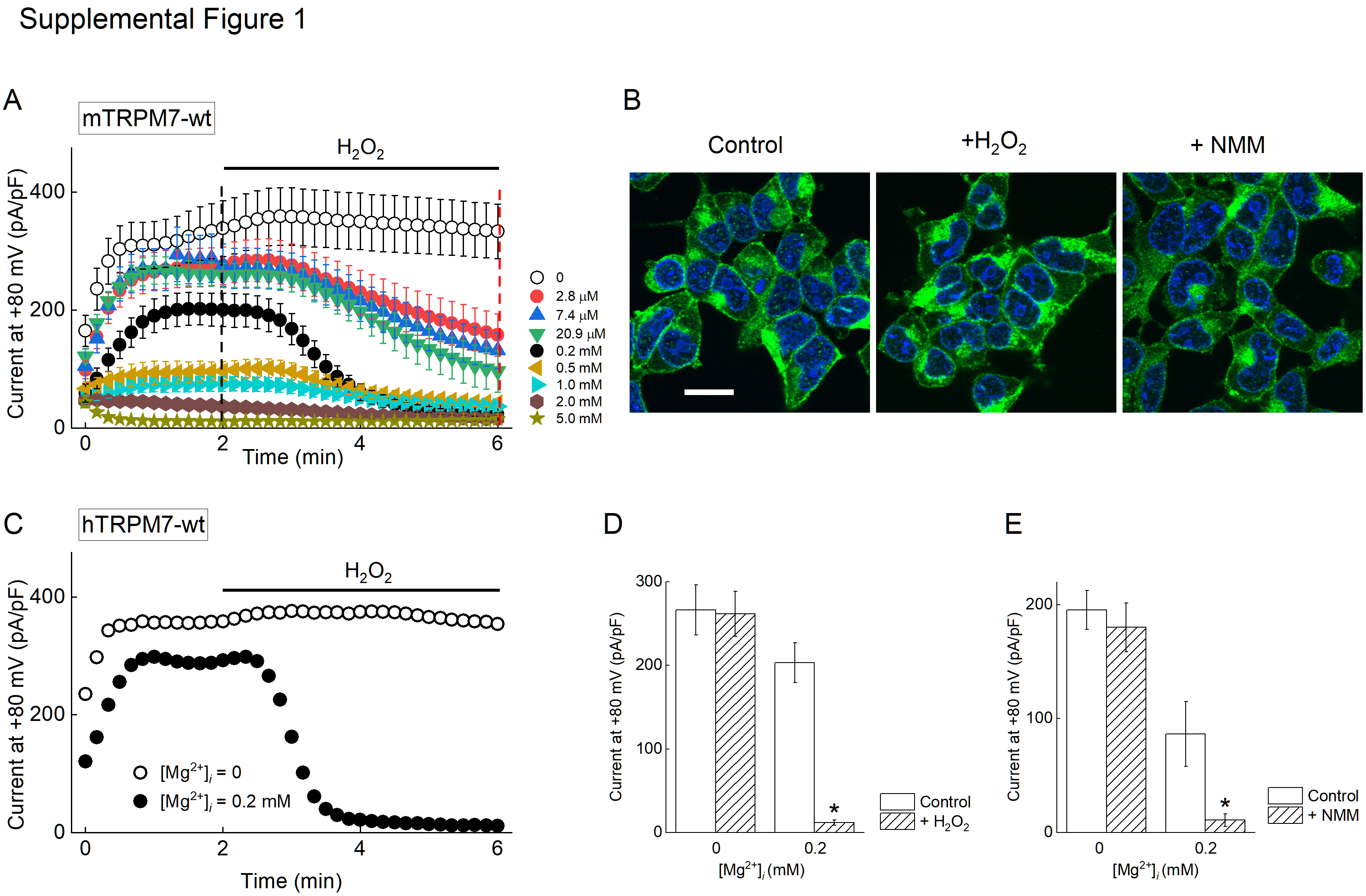
